# Distributed plasticity drives visual habituation learning in larval zebrafish

**DOI:** 10.1101/418178

**Authors:** Owen Randlett, Martin Haesemeyer, Greg Forkin, Hannah Shoenhard, Alexander F. Schier, Florian Engert, Michael Granato

**Affiliations:** Department of Cell and Developmental Biology, University of Pennsylvania Perelman School of Medicine, 1157 BRB II/III, 421 Curie Blvd., Philadelphia, PA 19104, USA; Department of Molecular and Cellular Biology, Harvard University, Cambridge, MA 02138, USA; Center for Brain Science, Harvard University, Cambridge, MA 02138, USA; Broad Institute of MIT and Harvard, Cambridge, MA 02142, USA; Harvard Stem Cell Institute, Cambridge, MA 02138, USA; Biozentrum, University of Basel, 4056 Basel, Switzerland

**Author notes:** Correspondence: O.R, F.E, M.G. Co-senior authors (Florian Engert, Michael Granato).

## Abstract

Habituation is a simple form of learning, where animals learn to reduce their responses to repeated innocuous stimuli. While habituation is simple in concept, its exact implementation in the vertebrate brain is not clear. It could occur via a single plasticity event at a singular site in the circuit, or alternatively via more complex strategies that combine multiple mechanisms at various processing stages and sites. Here, we use a visual habituation assay in larval zebrafish, where larvae habituate to sudden reductions in illumination (dark flashes). We find that 8 different components of this response habituate, including the probability of executing a response, its latency, and measures of its magnitude. Through behavioural analyses, we find that habituation of these different behavioural components occurs independently of each other and at different locations in the circuit. Further, we use genetic and pharmacological manipulations to show that habituation of different behavioural components are molecularly distinct. These results are consistent with a model by which visual habituation originates from the combination of multiple independent processes, which each act to adapt specific components of behaviour. This may allow animals to more specifically habituate behaviour based on stimulus context or internal state.

## Introduction

When presented with repeated but innocuous stimuli, animals habituate and thereby learn to ignore these potential distractions. This process of habituation is a form of learning that prevents an animal from wasting behavioural resources by responding to irrelevant stimuli. While habituation is referred to as the “simplest form of learning” [1], and appears to be conserved across all animals, the mechanisms underlying this process are still largely mysterious. Habituation is thought to occur via at least two temporally and molecularly distinct mechanisms, which lead to short-term memories that last for seconds to minutes, and long-term memories that last for hours or longer [1, 2]. Here we focus on long-term habituation, which, due to the extended time course, necessitates stable alterations to circuit properties [2–4].

In its simplest form, long-term habituation could result from a plasticity event at a single point in a circuit, such as a gain reduction at the sensory periphery, and studies of habituation in several species have focused on identifying the site and underlying mechanism of plasticity. These efforts have associated habituation with plasticity in upstream sensory-related brain areas, including: depression of the sensory-to-motorneuron synapse in the *Aplysia* Gill/Siphon withdrawal reflex [5, 6], depression of the sensory-to-interneuron synapse in *C. elegans* [7], and enhanced GABAergic inhibition in olfactory glomeruli in the *Drosophila* antennal lobe [8]. While there is substantial data confirming the importance of these upstream loci, it is possible that these individual sites are only one of many points in the circuit where plasticity is occurring. For example, the process of habituation may involve multiple plasticity events distributed throughout a neural circuit. Indeed, studies of short-term habituation in *C. elegans* indicate that multiple genetically separable mechanisms operate within a single habituation paradigm [9–11], supporting the model that multiple processes occur simultaneously during habituation. However, experimental evidence that multiple of such mechanisms also act simultaneously during long term habituation in vertebrates is currently lacking.

In the visual system, long-term habituation is especially challenging as it needs to be stimulus-specific, such that responses to some stimuli can be dampened without impairing vision in general. How and where in the circuit the necessary extraction and habituation of such stimulus features is executed is still largely unresolved. Inroads have been made in invertebrates, including the *Chasmagnathus* crab, which exhibits long-term habituation to a ‘visual danger signal’. This effect has been associated with decreased excitation of 3^rd^ order lobula plate neurons, but not in the upstream 2^nd^ order medulla neurons [12]. Examples in mammalian systems are usually more descriptive, and the precise underlying mechanisms are hard to unravel. Some progress has been made in mice, where it was shown that habituation occurs in response to specific orientations of moving gratings, and that this effect is linked to synaptic potentiation in the visual cortex [13]. Thus, it is tempting to speculate that more centrally located mechanisms of habituation are implemented in multi-functional systems like the vertebrate visual system.

In larval zebrafish, sudden reductions in illumination (dark flashes) induce a large angle turning manoeuvre called a “O-bend” [14]. Although the neural circuitry underlying the O-bend has not been well described, it is known to be retina-dependent [15]. At the reticulospinal level, the O-bend does not require the Mauthner neuron that drives the acoustic escape response [14], but does require the smaller and more ventromedially located reticulospinal neurons that also drive spontaneous turning behaviours (RoV3, MiV1, and MiV2) [16, 17]. Importantly, larval zebrafish exhibit protein synthesis-dependent long-term habituation to dark flashes, which, similar to memory formation in *Drosophila* and mammals, requires Neurofibromatosis 1 (Nf1) dependent cAMP- and RAS-mediated plasticity [18, 19]. Thus, dark flash habituation represents a tractable model for studying learning and memory in the vertebrate brain.

Here we show that the O-bend can be quantified by a defined set of kinematic parameters, and we find that these parameters are individually modifiable and therefore represent separately regulated components of behaviour. Importantly, we demonstrate that these components of the O-bend habituate concurrently, yet at different rates. Further, we show that habituation of these different components occurs via different molecular mechanisms, and that habituation can be selectively modulated by stimulus strength and the circadian rhythm. Combined, our results reveal that habituation, rather than originating from a single-point of plasticity, manifests from a surprisingly complex and distributed ensemble of processes that lead to modular habituation of specific behavioural components.

## Results

### High-throughput quantification of dark flash habituation

When exposed to a dark flash (a sudden transition to whole-field darkness), larval zebrafish execute an “O-bend” maneuver, characterized by a large angle body bend into an “O” shape followed by a swim forward in the new direction (Fig 1A, Movie S1) [14]. We developed a high-throughput behavioural setup that can track 600 larvae in individual wells, deliver visual and acoustic stimuli, and track stimulus responses at 560 Hz (Fig 1B, see Methods). This allows us to maintain individual larval identity throughout the experiment, to monitor behaviour over days, and to unambiguously classify stimulus responses using postural reconstruction of the bending axis of the larvae.

**Figure 1.**
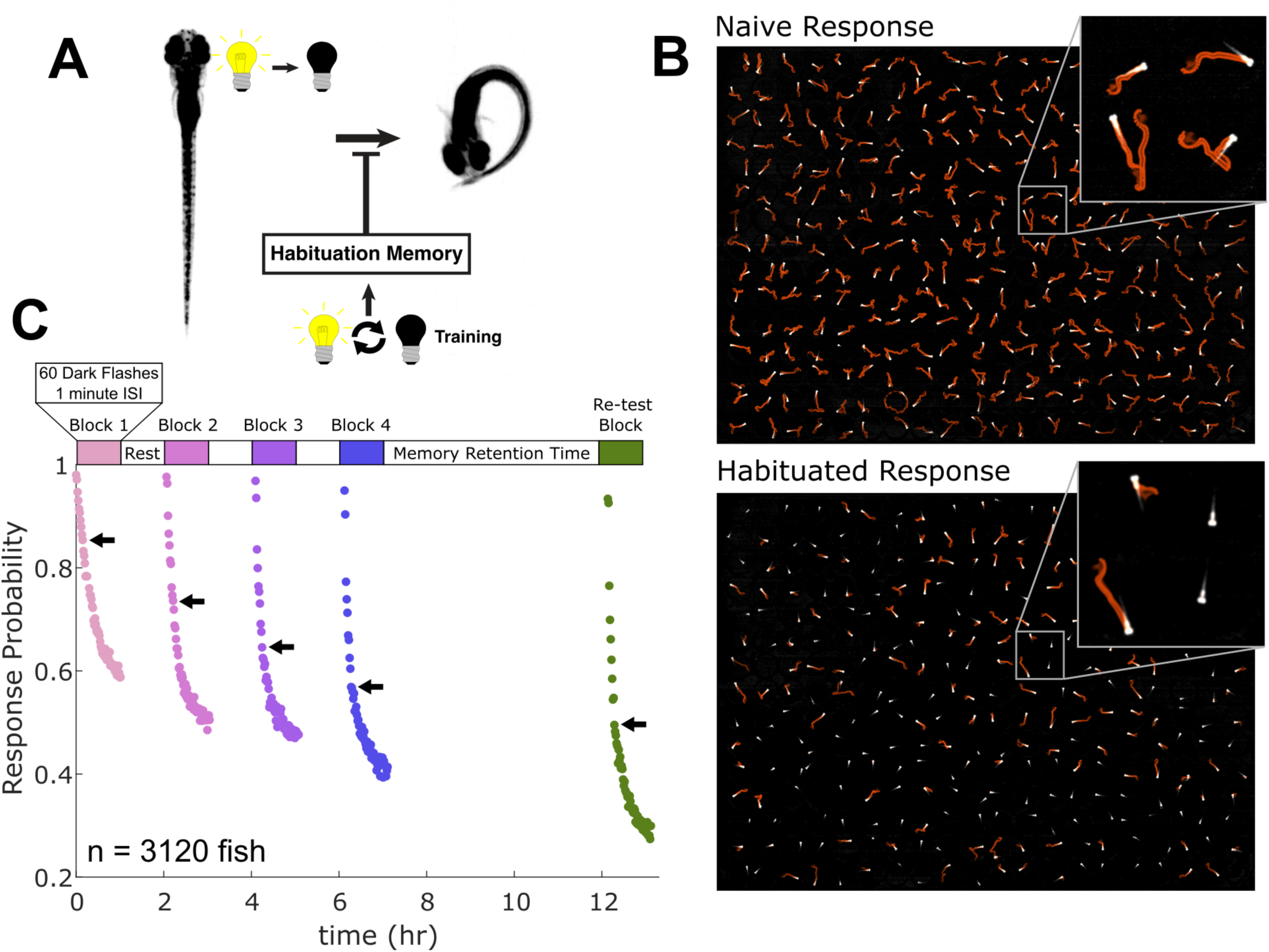
Habituation of the dark flash response. **A)** When stimulated with a dark flash, larval zebrafish execute a high-amplitude “O-bend” turn, which habituates with training [19]. **B)** Time projection images from a 0.9s recording of the same 300 larvae for the first flash (Naive Response), and the 240^th^ flash (Habituated Response). Motion is visible as orange streaks in the image tracing the path travelled by the larvae. Larvae that do not move are visible as stationary white larvae (insets). Larvae were recorded in 300-well plates, and images were background subtracted to remove the behaviour plates. **C)** The response probability across the population of larvae decreased both within the 60-flash training blocks, and successively across the 4 blocks of training. Each dot represents the proportion of larvae that respond to each stimulus, which are delivered at 1 minute interstimulus intervals (ISI). Memory was evident at the Re-test Block 5-hours after training, where larvae have not recovered to untrained levels (those in Block 1). Arrows = 10^th^ stimulus in each block.

Adapting an established dark flash habituation assay [18, 19], we developed a paradigm that consists of 4 training blocks of 60 dark flashes at 1 minute interstimulus intervals, with blocks separated by 62 minutes of rest. This spaced training paradigm induced habituation, which we quantified as the progressively decreasing probability of executing an O-bend (Fig 1B, C, Movie S2). Fitting curves to each block with an exponential function (Fig 2A) revealed that after each rest period, the response returned to near maximal levels, but then decreased more rapidly, and to lower values, during subsequent training blocks. This is consistent with previous observations of long-term habituation, which often manifests as a faster rate of re-habituation [1]. Memory retention was tested in a “Re-test” block 5 hours after training (Fig 1C), which revealed that O-bend behaviour did not recover to naive levels (those observed in the first block), but rather exhibited even greater reductions compared to the last block of training. Therefore, the habituation paradigm presented here induces memory that lasts for at least 5 hours.

**Figure 2.**
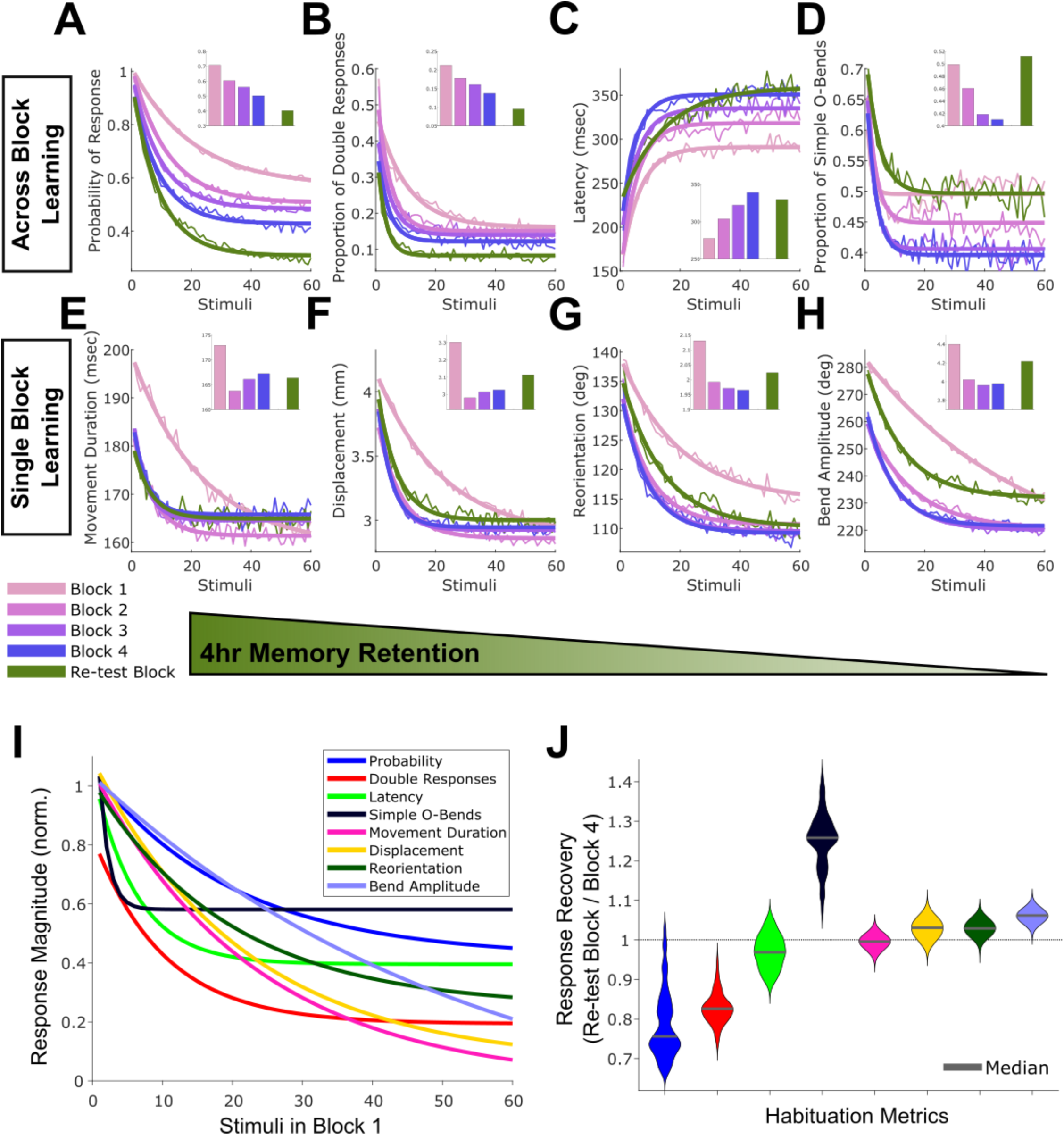
Habituation kinetics of different O-bend components. **A)** Exponential fits of habituation curves for each of the 4 training blocks and the re-test block plotted overlapping in time, colour coded by training block (thick line), thin line = raw data (mean across 3120 larvae), insets = mean response per block, for the probability of executing an O-bend, **B)** the proportion of larvae executing at least two O-bends, **C)** the latency from the stimulus to the initiation of the response (note that the latency values increase, indicating habituation), **D)** the proportion of larvae executing a simple O-bend, **E)** the duration of the movement, **F)** the displacement of the larvae, **G)** the reorientation angle achieved by the movement, and **H)** the maximal bend amplitude. **I)** Exponential fit curves for block one habituation performance plotted across components. Data was normalized such that initial response is equal to 1, and the minimal response observed in any flash is 0. **J)** Violin plots of the distributions of response recovery during the 5 hr. retention window, computed as the mean values across larvae for the trials in the Re-test block, divided by those in the last training block (block 4). Values greater than 1 reflect a recovery of the response.

Decreasing responsiveness is characteristic of habituation, but it is important to rule out fatigue as an alternative explanation. To that end, we monitored spontaneous movements (when no stimuli are delivered), the response to acoustic tap stimuli, and the ability of larvae to detect and respond to visual motion stimuli using the optomotor response (OMR) [20]. Specifically, we compared larvae that had undergone the 4-block training protocol with those that had not been trained. In each case, we did not detect reduced responses in the trained larvae, indicating that fatigue, or a generalization of habituation to other behaviours does not occur (Fig S1). In fact, rather than a fatigue induced reduction in motility, we observed small increases in displacement, turning rate and acoustic tap responses after training, indicating that dark flash habituation training may be slightly arousing to the animals. Importantly, as the OMR is also a retina-dependent behaviour, and as the OMR is unaffected by our training protocol, we conclude that habituation does not affect vision globally. This furthermore indicates that habituation is unlikely to occur at the general sensory neuron/photoreceptor-level, but rather selectively within the O-bend circuitry.

### Multiple O-bend components habituate

Habituation can be measured as a binary reduction in response (as above), or alternatively via decreases in the magnitude of the response. To further analyse this process, we asked how many other aspects of the response habituate, and subsequently how these might be related to one another. In total we identified 8 components of the O-bend that habituate (Fig 2A:H, Fig S2A):

**1) Probability** of responding to the stimulus, as discussed above. **2) Double Responses:** zebrafish larvae move in bouts, separated by periods of inactivity. By tracking behaviour over the full 1 second of the dark flash, we observed that many larvae do not execute a single O-bend, but rather execute multiple O-bends separated by at least 50 ms of inactivity (Movie S2). The proportion of larvae executing at least two responses habituates (Fig 2B). **3) Latency:** consistent with previous results [18], we observed that the latency from the stimulus to the response habituates, increasing by an average of 197 msec (first stimulus vs. last stimulus of block 4). Similar to components 1 and 2, we saw progressive accumulation of habituation across blocks, but retention after 5 hours was not as robust, though it still remained habituated compared to naive levels (Fig 2C). **4) Proportion of Simple O-bends:** in response to a dark flash, some larvae perform multiple high-amplitude bends in the same direction followed by a swim, as opposed to the classic O-bend that involves only one such bend before the swim. We term these “Compound O-bends” and “Simple O-bends,” respectively (Fig. S2B, Movie S2). Before training, 61% of the responses are Simple O-bends, which decreases to 41% with training(Fig 2D). We observed across-block habituation during training, but retention is poorer than components 1-3, and fully recovers to untrained levels after 5 hours.

We also observed habituation in O-bend components related to the kinematic magnitude of the movement, including its: **5) Duration**, **6) Displacement**, **7) Reorientation**, and **8) Bend Amplitude** (Fig 2E-H). Unlike components 1-4, habituation learning of these latter kinematic components occurred mostly during the first block, and did not decrease appreciably after block 2. Memory in these components was retained after 5 hours, with varying degrees of recovery. These results demonstrate that the O-bend rather than being an “all or nothing” response is actually composed of multiple behaviour components capable of adaptation during habituation. We noticed a significant degree of variability in learning and memory kinetics (Fig 2I, J), consistent with the idea that habituation of individual components rather than resulting redundantly from a single mechanism, instead result from multiple mechanisms with differential kinetics of adoption and decay.

### Habituation occurs at multiple circuit loci

To further explore the separability of the components of O-bend habituation, we took advantage of the spontaneous variability present in our dataset. Namely, while the majority of individual larvae habituate, there is considerable spread in the learning distributions (Fig 3A). We reasoned that if habituation occurs at a single circuit locus, then the learning performance of the different O-bend components would be correlated across larvae. In such a scenario, larvae would vary in their ability to habituate at this locus, but individual larvae would exhibit a consistent level of habituation across all behaviour components. Alternatively, if habituation of individual behaviour components occurs at distinct loci within the circuit, then learning performance would be independent of one another in any individual larva. Therefore, we would expect to see a lack of correlation in learning performance for these components across larvae.

**Figure 3.**
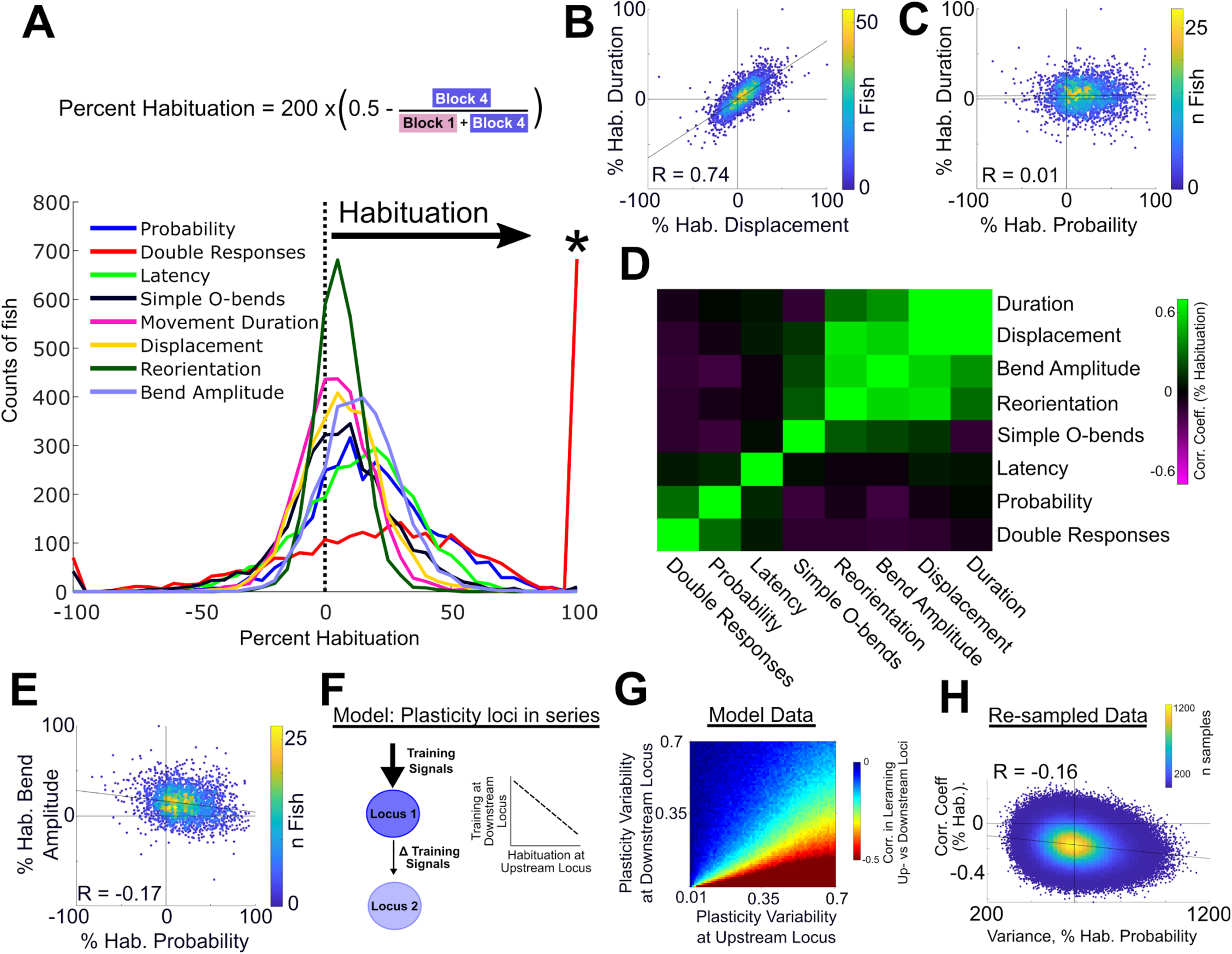
Habituation of different O-bend components occurs independently. **A)** Percent habituation histograms of individual larvae across the 8 O-bend components (n=3120 larvae). Asterisk marks the uptick in “Double Responses”, reflecting the individuals that show 100% habituation. **B)** Scatter plots and correlation coefficient comparing percent habituation for the “Displacement” and “Duration” components. Colormap reflects the density of points (R = 0.74, p<1×10^−10^, Spearman’s rho). **C)** Same analysis as B, revealing the “Probability” and “Duration” components are not correlated (R = 0.01, p = 0.48, Spearman’s rho). **D)** Hierarchically clustered correlation matrix (Spearman’s rho), comparing percent habituation of the O-bend components. **E)** Weak but significant anti-correlation observed between the “Bend Amplitude” and “Probability” components. **F)** Conceptual model for how anti-correlations of habituation performance could arise. If two plasticity loci exist in series and are habituating, in larvae where habituation at the upstream locus is stronger than average, this would result in less than average habituation training at the downstream node, and vice versa. **G)** Simulation results modeling plasticity at two loci that follow the kinetics of habituation for ‘Probability’, plus Gaussian noise. Training at the downstream locus depends on how much habituation occurs at the upstream node, resulting in anti-correlations in learning performance. The magnitude of the anti-correlations increases with learning variability at the upstream locus, and decreasing variability at the downstream locus (n = 10000 runs per comparison). **H)** Re-sampling of the original 3200 larvae into random 100-larvae subsets over 2000000 iterations. Variance in percent habituation for “Probability” scales with the magnitude of the anti-correlation between percent habituation for “Probability” and “Bend Amplitude (p < 1×10^−10^, Spearman’s rho).

Our analysis indicates that both scenarios occur. We observed strong positive correlations between some components, such as “Displacement” and “Movement Duration” (Fig 3B). These two components also show similar learning and retention kinetics (Fig 2I, J), further supporting the idea that they habituate through a single mechanism. However, other components, such as “Probability” and “Movement Duration” showed no correlation (Fig 3C), demonstrating that the capacity of a larva to learn to respond less frequently is uncoupled from its ability to learn to respond with a shorter movement. Analysis of all pairwise comparisons (Fig 3D) revealed that learning was largely correlated across the kinematic components (5-8). “Probability” and “Double Responses” are also correlated, while “Latency” and “The Proportion of Simple O-bends” are not strongly correlated to any other group. This leads us to conclude that dark flash habituation depends on plasticity exerted at four or more distinct loci.

We also observed weak negative correlations between habituation of some O-bend components, most prominently between “Probability” and “Bend Amplitude” (Fig 3E). One possible explanation for such subtle anti-correlations might be due to the circuit architecture of habituation. For example, if different plasticity loci in the circuit operate in parallel, we would expect to observe no correlation in learning performance for their respective components. Alternatively, if the loci are arranged in series, habituation at the upstream locus will reduce the amount of training signal that reaches the downstream locus. Since habituation results from repeated stimulation, this would result in a negative relationship between upstream plasticity and downstream training (Fig 3F). To demonstrate that this can occur, we modelled two habituating neurons connected in series. Both neurons acted as habituation loci with the same habituation kinetics, with random noise added to simulate learning variability. This simulation confirmed that anti-correlated distributions manifest from such an architecture, and that the magnitude of the anti-correlation increases with variability at the upstream plasticity locus (Fig 3G). If we assume from a sensorimotor perspective that the locus that habituates “Probability” is upstream of “Bend Amplitude”, then this simple model predicts that the magnitude of the anti-correlations in habituation for “Probability” and “Bend Amplitude” would increase with the variability for “Probability”. Using iterative sub-sampling of groups of 100 larvae, we observed that indeed, the magnitude of the anti-correlations increases along with the variance in percent habituation for “Probability” (Fig 3H). Combined, these results support a model by which habituation results from distributed effects spread across multiple circuit loci, some of which operate in parallel in the circuit (no correlation in learning performance across larvae), while others operate in series, resulting in negative correlations.

### Habituation of different O-bend components are molecularly separable

Although our results indicate that multiple sites in the O-bend circuit exhibit plasticity independently during dark flash habituation, it is unclear if these distinct events use distinct molecular pathways. If different molecular pathways operate, then it should be possible to identify manipulations that differentially affect different O-bend components. To test this, we analyzed *neurofibromatosis 1* (*nf1*) mutants, which fail to habituate the latency of their dark flash responses [18]. When we analyzed habituation in *nf1* mutants, we found that not all components are equally affected (Fig 4A). In fact, while learning performance is strongly inhibited for “Latency” (Fig 4B), learning performance is indistinguishable from controls for “Displacement” (Fig 4C), strongly suggesting that individual components of dark flash habituation are regulated via distinct molecular mechanisms. To further generalize these findings, we next performed a set of pharmacological manipulations. Due to their previously identified roles in zebrafish behavioural plasticity and habituation, we tested antipsychotic drugs that act as antagonists of the dopamine and serotonin systems [19, 21, 22]. Specifically, treatment with Haloperidol, a dopamine D2 receptor antagonist, had a wide range of effects on habituation (Fig 4D). Remarkably, these effects include oppositely signed effects for different O-bend components, such as increased habituation for “Latency” (Fig 4E) and decreased habituation for “Bend Amplitude” (Fig 4F). Similarly, treatment with Pimozide and Clozapine, which also antagonize the dopamine D2 receptor, had separable effects across different behaviour components (Fig 4G, H). These pharmacological experiments confirm that habituation of different O-bend components occurs via different molecular mechanisms.

**Figure 4.**
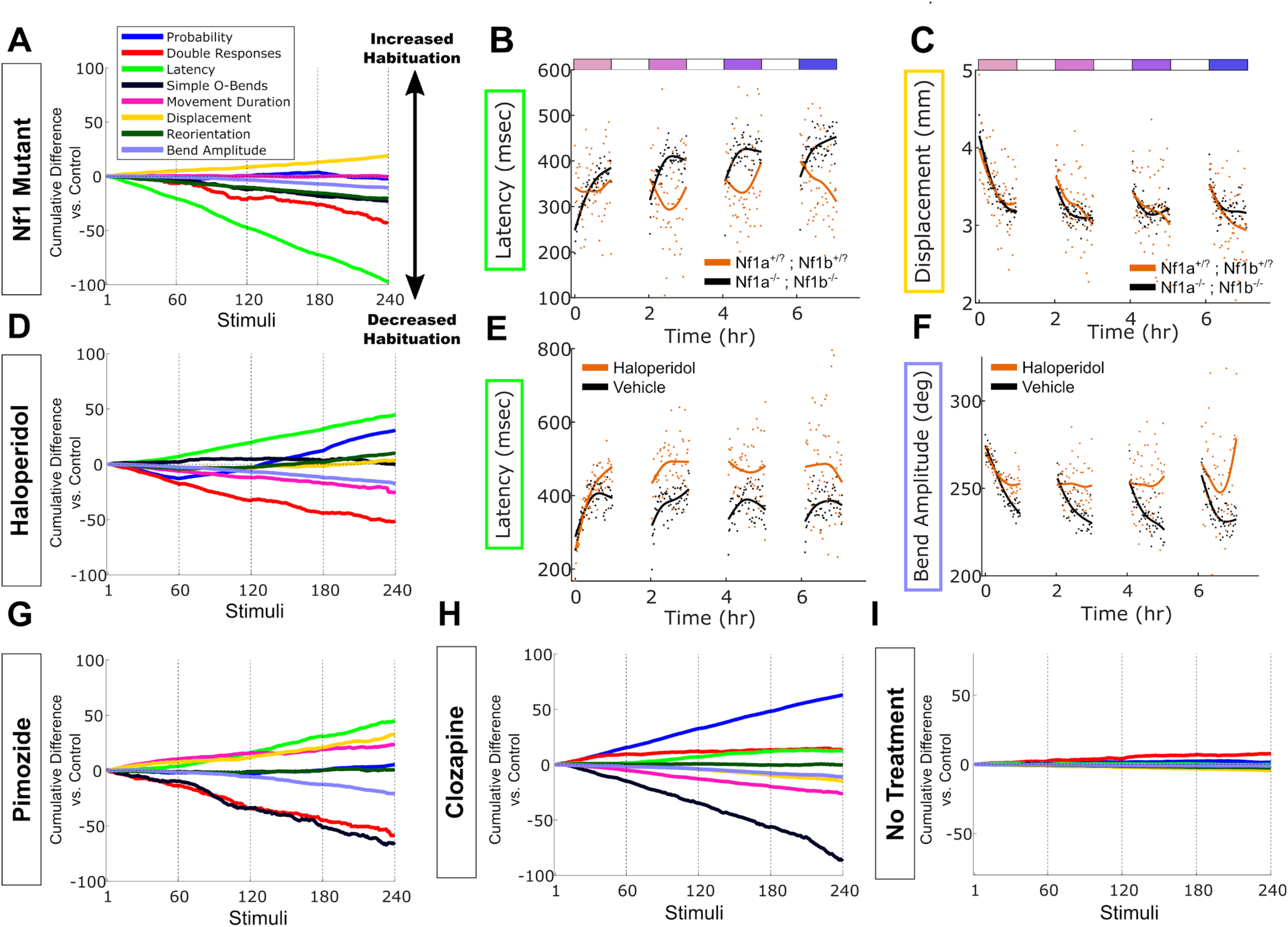
Habituation of different O-bend components are genetically and pharmacologically separable. **A)** Cumulative plot of the differences in habituation rate for the O-bend components. These plots display the cumulative differences in the mean response of *nf1a;nf1b* double homozygous mutants (n=29 larvae) compared to sibling controls (n = 444 larvae). Difference from 0 reflects divergence in response across the 240 dark flash stimuli in the 4 training blocks, with negative values reflecting a failure to habituate. Mutants fail to habituate some components, most profoundly for the “Latency” metric. **B)** Raw data (dots = mean across larvae for each stimulus) and smoothing spline fits (solid lines), demonstrating that *nf1a;nf1b* double mutants fail to habituate when measuring “Latency” (note that increasing latencies indicate habituation, p = 2×10^−12^, Wilcoxon signed rank test). **C)** *nf1a;nf1b* double mutants habituate normally when measuring “Displacement” (p = 0.92, Wilcoxon signed rank test). **D)** Cumulative difference plots for treatment with Haloperidol (10 μM, n = 80 larvae) vs. vehicle controls (0.1% DMSO, n = 140 larvae). **E)** Treatment with haloperidol increases habituation performance when measuring “Latency” (p = 9 × 10^−26^, Wilcoxon signed rank test), and **F)** decreases habituation when measuring “Bend Amplitude” (p = 9 × 10^−20^, Wilcoxon signed rank test). **G-J)** Cumulative difference plots comparing 0.1% DMSO vehicle controls and after treatment with **G)** Pimozide (1μM, n = 20 treated larvae, n = 60 controls), **H)** Clozapine (10 μM, n = 120 treated larvae, n = 160 controls), and **I)** comparing wild type larvae in the same experiment that are given no treatments as a negative control (n= 150 larvae, both groups)

### Habituation of different O-bend components are separably modulated by stimulus strength and the circadian rhythm

While we have shown that habituation of different O-bend components is independent, it was unclear what utility such independence might serve. We reasoned that having a modular system might allow an animal to adapt its behaviour with more flexibility or specificity in different contexts. To test this, we first asked how habituation rates change when the stimulus is weakened. Instead of delivering a full dark flash, we decreased the illumination by only 80%. This weakened stimulus is still strong enough to reliably elicit O-bends [14], and causes the larvae to habituate more rapidly (Fig 5A-C). However, the effect was selective for the “Probability” component, while other components, including “Bend Amplitude”, showed much less modulation. This indicates that the nature of the stimulus can alter the habituation rate of different behavioural components in different ways.

**Figure 5.**
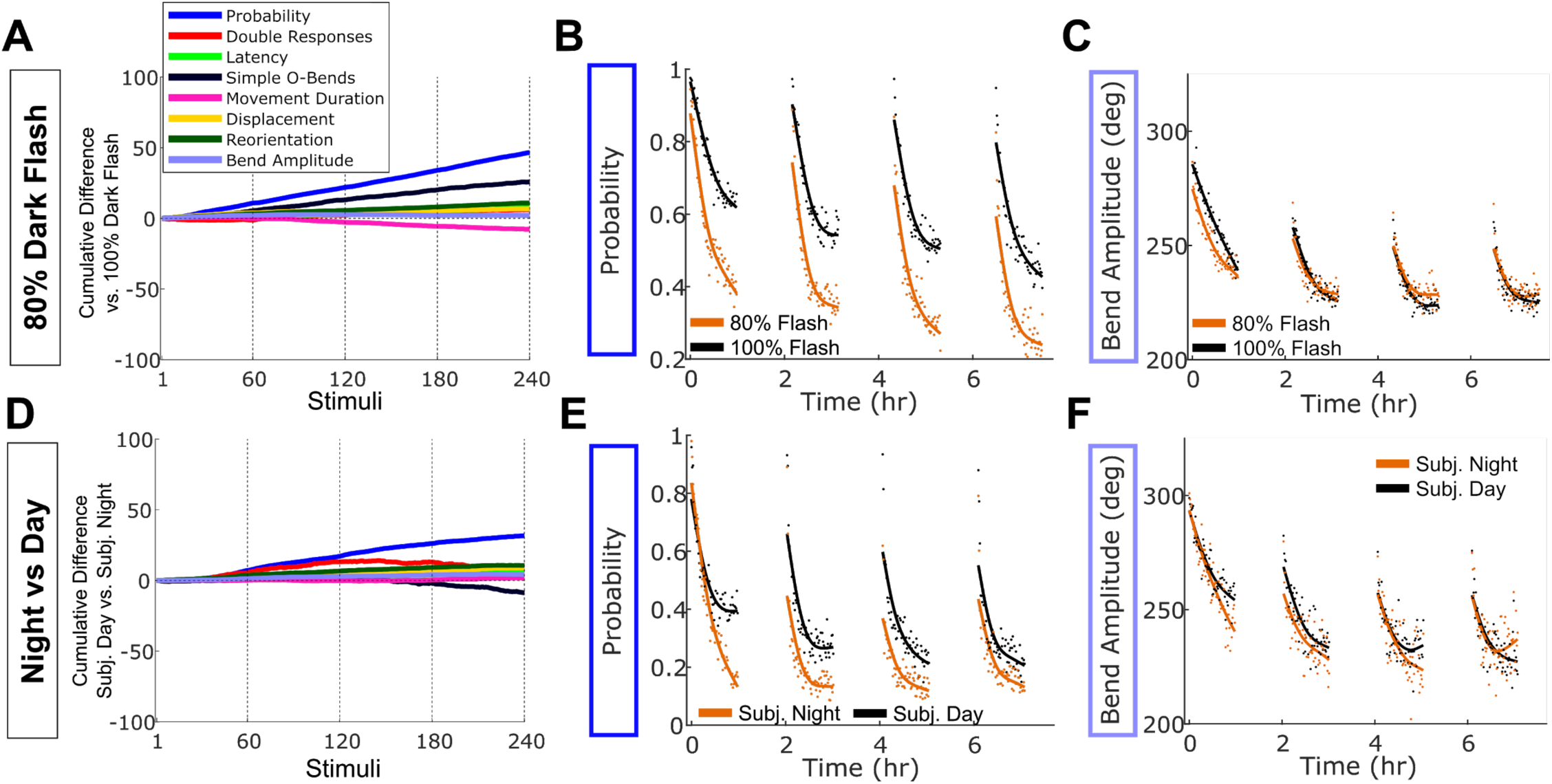
Habituation of the “Probability” component is modulated by stimulus strength and circadian phase. **A)** Cumulative difference plots comparing larvae given an 80% dark flash (light intensity transitions from 100% to 20%), with larvae given a normal 100% dark flash (n = 300 larvae, both groups). **B)** Raw data (dots = mean across larvae for each stimulus) and smoothing spline fits (solid lines), showing how larvae habituate more rapidly to the weaker stimulus for the “Probability” component (p = 4 × 10^−41^, Wilcoxon signed rank test), while **C)** habituating the “Bend Amplitude” more similarly to controls, though a significant difference was detected in the distributions (p = 0.02, Wilcoxon signed rank test) **D)** Cumulative difference plots comparing larvae in the subjective night phase of the circadian cycle with those in the subjective day (n = 150 larvae, both groups). **E)** Larvae in the subjective night phase show increased habituation performance for the “Probability” component (p = 9× 10^−39^, Wilcoxon signed rank test), and **F)** much weaker effects on “Bend Amplitude” (p = 3 × 10^−7^, Wilcoxon signed rank test)

Finally, to further test our hypothesis that a modular system enables animals to adapt habituation behaviour in a more context dependent manner, we compared habituation rates during different phases of the circadian rhythm. The circadian phase modulates the endogenous arousal level of zebrafish larvae [23], and may also alter the salience of a dark flash since darkness is an expected condition at night. Specifically, we raised a subset of larvae on a reversed night/day light cycle and subjected them to our habituation assay together with their normally raised siblings. By testing the behaviour of larvae during either their subjective day, or their subjective night, we found that there was a circadian influence on habituation (Fig 5D-F). Similar to the effect of a weakened stimulus, we identified selective effects on the habituation rate of the “Probability” component, which showed significantly stronger habituation during the night phase. This indicates that modularity in habituation is exploited endogenously by the larvae, where stimulus strength and the circadian rhythm have similar and specific effects on the habituation rate of different O-bend components. This could allow the larvae to more rapidly cease responding during the night, or to weaker dark flashes, while continuing to adapt kinematic-related components at the normal rate. Thus, modularity in habituation can allow for the adaptation of specific behavioural components based on both the context of the animal and the stimulus.

## Discussion

The experiments presented here quantify behaviour across thousands of larval zebrafish and demonstrate that dark flash habituation occurs via multiple plasticity events, where each of these events acts to suppress a specific component of behaviour. These different plasticity events manifest in differential kinetics of learning and forgetting and a lack of correlated learning across behavioural components. Particularly surprising was the separability of “Probability” and “Latency”, since in the simplest model, “Latency” is a direct function of “Probability” (as in a Poisson process, similar to [24]). Since the brain can adapt these two aspects of behaviour separately, the decision of whether to respond to a stimulus appears to be uncoupled from the decision of when to respond.

Our experiments indicate that the brain not only implements plasticity in multiple circuit loci, but also does this via multiple molecular mechanisms. We found that some components of habituation require Nf1, while others do not. Furthermore, our experiments with antipsychotic drugs indicate that some events involve signalling through dopamine and/or serotonin receptors, while others do not. This clearly opens the path for a whole series of detailed investigations on the precise nature and mechanistic role of these pathways that will be the subject of future studies. However, as these drugs all antagonize the dopamine D2 receptor, and all increase habituation for “Latency”, it is likely that dopamine signalling negatively regulates this aspect of habituation. The effects of opposite sign that a single drug can have across different components suggest that the same molecular pathways are capable of influencing different plasticity events in oppositely signed manners. Alternatively, these effects may relate to the promiscuous nature of these drugs, which can affect many targets [25]. Considering that these drugs are used to treat schizophrenia, and that there are well-established connections between habituation and schizophrenia (as well as other psychiatric disorders including autism [26, 27]), our approach that allows to disambiguate specific behavioural components in a high-throughput assay may have important relevance for translational approaches. For example, it might aid efforts aimed at identifying more selective therapeutic compounds that share targets with known beneficial pharmaceuticals, but that act with greater molecular and behaviour-modifying precision.

Habituation of different O-bend components may result from plasticity at different synapses within the same neurons, but a more parsimonious mechanism would involve different neurons that are part of parallel or serial pathways within the circuit. The negative correlations in habituation performance that we observed in some components also support a model where plasticity occurs in different neurons that are arranged in sequence within a sensory motor path. This can be explained by a simple model, where individual variations in plasticity at upstream neurons result in variable levels of activation and thereby variable opportunity for habituation at downstream neurons. The physical location of plasticity sites remains to be determined. However, it is tempting to speculate that plasticity regulating the release of the O-bend response, such as “Probability” and “Latency”, might exist more towards the earlier sensory-related parts of the circuit, while regulation of kinematic parameters might occur downstream towards the motor circuitry in the hindbrain or spinal cord.

In light of recent work in *C. elegans*, where multiple genetically separable mechanisms have been shown to underlie short-term habituation [9–11], we propose that such modularity is a conserved feature of habituation. Thus, to accurately identify and characterize the possible neural implementation of habituation, it is important to consider behaviour not as a single output, but rather as a combination of multiple independent modules. Why might the zebrafish brain have evolved such a seemingly complex strategy to habituate? Perhaps plasticity to repeated stimulation is simply a pervasive adaptation at many synapses in a circuit, and we can observe these multiple effects when analyzing behaviour in a multi-component manner. Alternatively, approaching habituation in a modular way would allow for greater flexibility. This would allow for more specific adaptations than a simple global reduction in responses, perhaps tuned based on brain state, stimulus, or environmental context. Indeed, we found that habituation of “Probability” is modulated by the circadian rhythm and dark flash intensity, while other O-bend components are not. This demonstrates that the modularity of habituation manifests endogenously, and acts to tune the habituation rate of different components of behaviour based on the context of the animal. Thus, our results reveal that the strategies taken by even relatively simple larval zebrafish brains to habituate require a surprisingly complex coordination of different plasticity events distributed across the circuit.

**Figure S1.**
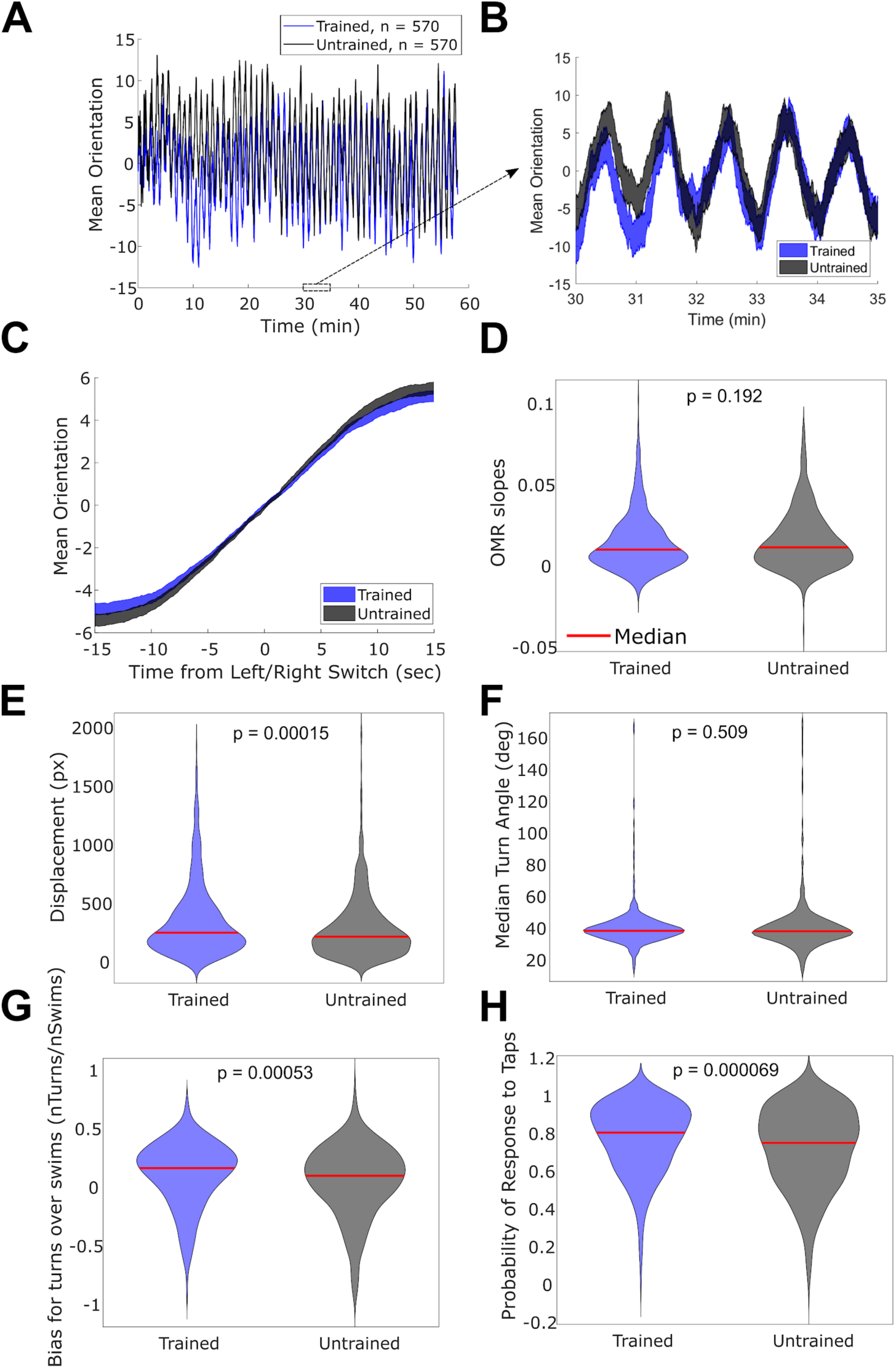
(related to Figure 1). Dark flash habituation does not generalize to other behaviours. **A)** The optomotor response (OMR) is induced by an approximate sine wave, alternatively traveling in leftward and rightward directions, switching directions every 30 seconds. This induces an oscillating pattern in the average orientation of larvae, which is **B)** locked to the directional transitions at 30s intervals, and similar in amplitude in trained and untrained larvae (mean across larvae +/- SEM, n = 570 trained and 570 untrained larvae, in all panels). **C)** These oscillating patterns can be recomposed and averaged into a single 30 second long period, where OMR performance is indicated by the positive slope of the line. **D)** These slopes are not significantly different when comparing trained and untrained larvae. Data is shown as violin plots showing the smoothed distributions, and tested for statistical significance using the Wilcoxon rank sum test (panels D-H). **E)** When not delivering any stimuli over a 30 minute period, the spontaneous displacement of the larvae is not suppressed by training, but rather slightly larger in the trained group. **F)** The median turning angle of the larvae is not significantly different, and **G)** there is a slight but significantly higher bias for swims over turns after training, indicating that the larvae readily perform turns after training. **H)** The response rate to a different stimulus modality was tested by delivering 30 acoustic taps to the larvae, which induce C-start escape responses. Trained larvae were slightly but significantly more likely to respond to these taps, indicating that dark flash habituation training does not generalize across these stimulus modalities.

**Figure S2.**
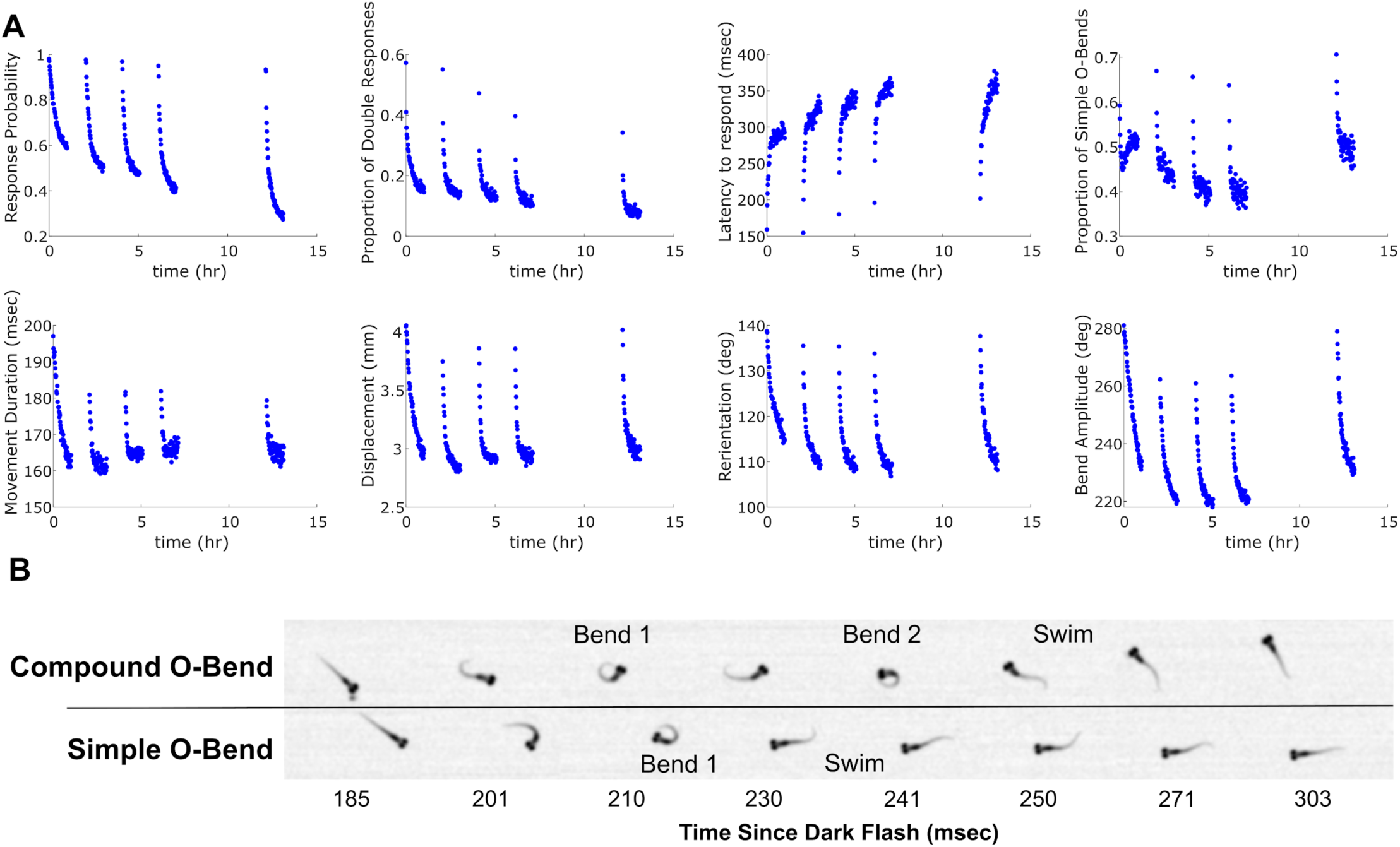
(related to Figure 1). Habituation curves across behaviour components. **A)** Habituation is evident in 8 different components of the response, each dot represents the mean response to each stimulus (n=3120 larvae). **B)** A “Compound O-bend” is defined as a movement where a large angle bend (Bend 1) is followed by a period of partial relaxation of the bend, before performing a second large angle bend (Bend 2). Finally, the larva fully relaxes and swims forward. In contrast, a “Simple O-bend” is characterized by a single large angle bend (Bend 1), which fully relaxes and the larva swims forward.

**Movie S1**. The O-bend response.

The same larva performing an O-bend to three different dark flashes. The O-bend is characterized by an initial high-amplitude “O”-shaped bend, followed by a swim in the new heading direction. The movies are synchronized to the beginning of the movement, and last for 197 msec.

**Movie S2.** Habituation of the O-bend.

Movies of the same 300 larvae comparing the response to the first dark flash (Naive Response, left), and the 240^th^ flash (Habituated Response, right). Larvae are recorded at 560hz in 300-well plates. The images are background subtracted so that the wells are not visible and the larvae appear white. The cumulative path taken by the larvae becomes visible as orange streaks in the movies to highlight the responses.

**Movie S3.** Compound vs Simple O-bends, and double responses to a single dark flash.

Recordings of two larvae, cropped from movies of the 300-well behaviour plates. Time is displayed relative to light-offset, which remains off for the entire movie. The larva in the bottom well performs a Simple O-bend, characterized by a single high-amplitude bend, followed by a swim forward in the new heading direction. The larva in the top well performs a Compound O-Bend, characterized by two separate high-amplitude bends in the same direction, followed by a swim forward. After a period of immobility, this larva then performs a second response, this time a Simple O-bend. The larva on the bottom also moves a second time, but performs a “swim” without any characteristic high-amplitude O-bend.

## Methods

### Experimental model and subject details

Experiments were conducted on 5-6 dpf larval zebrafish (*Danio rerio*, TLF strain) raised in E3 medium at 29°C on a 14:10 hr light cycle. For the circadian experiments (Fig. 4 J-L), larvae tested in the subjective day were raised on a 9am-ON, 11pm-OFF light cycle, while larvae tested in the subjective night were raised on an 11am-OFF, 9am-ON light cycle, and both groups were tested beginning at ∼12:30pm. Breeding adult zebrafish were maintained at 28°C. Behavioural assays on larvae carrying mutations for Nf1a^p301^ (ZDB-ALT-130528-1) and Nf1b^p303^ (ZDB-ALT-130528-3) were conducted blind to genotype. Subsequent genotyping was performed with the KASP method with proprietary primer sequences (LGC Genomics). This method was validated using previously described PCR genotyping [28]. All animal protocols were approved by the University of Pennsylvania Institutional Animal Care and Use Committee (IACUC).

### Behaviour recording

Larval behaviour was recorded in multiwell plates fabricated from 6.3mm thick clear acrylic sheets (US Plastics). The acrylic was cut with a laser cutter into 8mm diameter wells with a volume of ∼ 300 uL, arranged in a 20×15 grid for a total 300-well plate. 3.2mm white acrylic (US Plastics) was bonded to the cut wells (SciGrip 4), acting as both the bottom of the plate and the light diffuser. To minimize evaporation and to maintain a consistent ∼28°C temperature in the behaviour wells, the 300-well plate was placed under a 29-31°C water bath that acted as a heated lid for the plate.

Larvae were illuminated from below with IR LEDs (890nm, Vsiahy.com part number TSHF5410) driven by a 1A Buckpuck driver (Luxdrive). Images were recorded from above with EoSens 4CXP Monochrome Camera (Mikroton), an 85mm 1.8 AF D lens (Nikon) with a IR long-pass filter (LP780-62, Midwest optics), and a Cyton Quad Channel CoaXPress Frame Grabber (Bitflow). The camera was triggered at 560hz using a Teensy 3.2 microcontroller (PJRC).

Due to the symmetrical design of the behavioural assay (1 hr training blocks, 1 hr rest between blocks), we were able to double the throughput of the rig by alternatingly imaging two separate 300-well plates during the experiment. The larvae in plate 1 were recorded during training, and during the rest period the camera view was switched to plate 2, which was trained and recorded during the rest period for plate 1 (and vice versa). Therefore, the experimental time for the first and second plates are offset by one hour. Switching the camera views was done by placing the camera at a 90 degree angle above the behaviour plates and using two 4’’ x 5’’ 45-degree incidence hot mirrors (43-958, Edmund Optics) to direct the camera view towards the two behaviour plates. The mirrors were attached to Nema 17 stepper motors (ROB-09238, Sparkfun), driven by an EasyDriver (ROB-12779, Sparkfun), a Teensy 3.2 microcontroller (PJRC), and the Multi-fish-tracker software. In this way they could be rotated in and out of place to view each plate. Light cross talk between the behaviour plates was minimized using blackout hardboard (TB4, Thorlabs).

For visual stimuli, we made a rectangular ring of 115 RGB LEDs (WS2812B 5050 RGB LED Strip 1M 144LED/M, ebay.com) to border the 300-well plate, diffused by 3.2mm white acrylic (US Plastics). LEDs were controlled using a Teensy 3.2 microcontroller and the fastLED Animation library (http://fastled.io/). The LEDs were set to white color with a brightness value of 50, yielding an intensity of approximately 130uW/cm^2^ at the behaviour plate. During a dark flash, the LEDs were turned off for 1 second and video of larval responses was recorded at 560 Hz. After this, the light intensity increased linearly to the original brightness over 20 seconds. To induce the optomotor response, an approximate sign wave was generated by illuminating every 8th LED along the top and bottom of the plate. The position of the illuminated LED was progressively shifted down the strip by ramping down the intensity of the illuminated LED, while ramping up the intensity of the adjacent LED. In this way, the motion stimuli were translated in the leftward and rightward directions relative to the plate. The direction of motion was switched every 30 seconds, for a total testing period of 1 hour. The orientation of the zebrafish larvae was tracked online using the Multi-fish-tracker (see below) at 28 Hz and was used to quantify the optomotor response, which follows the direction of perceived motion. Acoustic tap stimuli were delivered using a Solenoid (ROB-10391, Sparkfun) that delivered a single tap to the top of the water bath and induced acoustic escape responses.

### Multi-fish-tracker

The code to track individual zebrafish larvae in a multi-well format was custom written in C# (Microsoft, USA) using Intel’s integrated performance primitives (IPP, Intel, USA) for fast image processing. Specifically, a running average background was kept for each plate that was updated with an exponential decay time of 2 minutes. This was done to flexibly adapt to different lighting conditions. The plate was subsequently divided into two sections, which were tracked on separate threads to increase throughput. Individual wells were identified using user-defined masks. The background was subtracted from each image and the resulting absolute difference was thresholded. Subsequently, the biggest object in each well, physically close to a previously identified larval position, was designated as the larval object, and relevant parameters such as position and heading angle were extracted using image moments. At baseline, tracking was performed at 28 Hz. For one second after each dark flash (or tap), all frames at the full camera frame-rate of 560 Hz were written to disk, for detailed offline kinematic analysis of behaviour.

### Offline video tracking

Offline tracking on recorded videos was performed in Matlab (Mathworks). The image was background subtracted and thresholded to identify the centroid of the larvae in each well. The background subtracted image was convolved with a head-sized filter, and the maximum intensity pixel was used to identify the head coordinate. Then, using the search direction vector defined by the head-to-centroid direction, 8 points were placed along brightest points on the larvae. The head coordinate was used to calculate speed and displacement of the larvae, the head-to-centroid vector was used to calculate the heading orientation of the larvae, and the cumulative angle between the tail points was used to calculate the bend amplitude of the larvae.

### Behavioural quantification

Analyses of larval behaviour and statistical analyses were performed in Matlab (Mathworks). For each dark flash or tap stimulus, the offline tracked videos were used to score behaviour during the 1 second of recorded video. O-bends in response to dark flashes were identified as movement events that had a bend amplitude greater than 3 radians (172 degrees). C-bend responses to taps were identified with a bend amplitude greater than 1 radian. Compound O-bends (Fig S2B) were classified as O-bend events which had at least two local maxima in the bend amplitude trace during the initial bend before the bend amplitude trace crossed 0.

The proportion of the larval population that performed an O-bend at each dark flash was used to quantify habituation performance for “Probability” (Fig 1C, 2A). To generate habituation curves for the other behavioural components (Fig, 2B-I, S2A), larvae that did not perform an O-bend for a given stimulus were excluded from the analysis at that stimulus. To fit exponential curves to each training block of 60 flashes (*x*), we used Matlab’s ‘fit’ function, with a ‘fittype’ formula of:

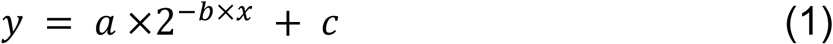

To quantify the recovery of the response in the population (Fig 2J), we averaged the response at each flash in Block 4 and the Re-test block across all larvae, and divided the Re-test block responses by the responses in Block 4. These distributions were plotted with ‘violin.m’ [29], with a bandwidth of 0.15.

Percent habituation was calculated for each larva as the decreased mean responsiveness for the 60 flashes in training Block 4, relative to the mean response for the 60 flashes training Block 1, using formula (2). To make the distributions comparable across the different components of behaviour, the minimum observed mean value across all larvae was subtracted from the block 1 and block 4 mean responses. This ensures that the responses can scale all the way to 0 regardless of behaviour component -for example “Curvature”, which by our definition of O-bends, must be at minimum 3 radians. Except when calculating for the “Probability” behaviour component, larvae that did not perform an O-bend for a given stimulus were excluded from the analysis at that stimulus.

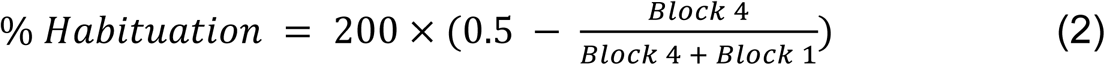

Analysis of stimulus-free swimming behaviour (Fig S1E-G) was done using the online tracked larval coordinates from the Multi-fish-tracker. This was done for a 30 min period beginning one hour after the fourth dark flash habituation training block. Analysis of the optomotor response (OMR, Fig S1A-D), was done using the heading angle from the Multi-fish-tracker for one hour, beginning 3 hours after the fourth dark flash habituation training block. To calculate OMR performance, we isolated each 30-second period where the larvae were either being stimulated with leftward or rightward motion. We isolated the left-right component of the orientation by calculating the arcsin of the sin of the larval orientation. We then reflected the traces in time during the rightward stimulus presentation, such that each 30-second period would have an increasing slope if the larva were to reorient to follow the direction of motion (Fig S2C). All the 30 second periods were then averaged for each larva, and these averaged traces were fit with linear regression using Matlab’s ‘polyfit’ function. The slope of this fit (OMR slope) was taken as the measure for OMR performance for each larva (Fig S2D).

To calculate the cumulative difference in habituation performance for the Nf1 mutants, pharmacological treatments, 80% flash, and the circadian experiments (Fig 4, 5), we calculated the average response across larvae at each dark flash. This was done for the treatment and control groups, yielding a mean vector for each group. These two vectors were normalized by dividing them by the initial response to the first flash for each group, and they were then subtracted, yielding the a mean difference vector between stimulus and controls. If the two groups are habituating similarly, then these difference vectors will have a mean of approximately 0, and thus the cumulative distribution would remain near 0. However, treatments that affect habituation will show strong increasing or decreasing cumulative distributions, reflecting increased or decreased habituation performance throughout training, respectively. We confirmed this by comparing larvae in even and odd numbered wells, which showed little divergence from 0 in the cumulative distributions (Fig 4I). Reported p-values were calculated by subtracting the mean control response vector from each treated larva’s response vector, and by taking the mean of this difference vector. A difference from 0-mean in the treated population was tested for using the Wilcoxon signed rank test (‘signrank’ in Matlab).

### Modeling

We modeled two habituation loci acting in series in Matlab. We began with the upstream locus, which follows learning kinetics of the exponential fit for the first training block of 60 flashes when measuring “Probability” (Fig 2A, Block 1, Equation (1)). In each model run, gaussian noise was added to the coefficients of the fit, resulting in variable habituation curves. Each learning curve was normalized such that the maximum value is equal to 1. The downstream locus follows the same learning kinetics, but the learning curve was truncated based on how much habituation occurred at the upstream locus. This was measured as the area under the habituation curve at the upstream node. If little habituation occurs, this will be close to 60, for the full 60 flashes. However, if habituation is profound this will approach 1. Therefore, the opportunity for habituation at the downstream node is negatively dependent on habituation performance at the upstream node. The variability in habituation across runs (axes in Fig 3G) was controlled by varying the standard deviation of the gaussian distribution from which the noise added to the coefficients was derived. Learning performance was defined as in Equation (2), replacing Block 1 with the initial value of the habituation curve, and Block 4 with the final value of the curve. For each value of habituation variability (‘sigma’) at each locus, 10000 iterations of the model were run. The correlation in learning performance at each node across model runs (Spearman’s rho) was calculated using Matlab’s ‘corr’ function.

### Pharmacology

Stock solutions of 100uM Haloperidol (H1512, Sigma), Pimozide (P1793, Sigma) and Clozapine (C6305, Sigma) were prepared in DMSO. 10x solutions in 1% DMSO in E3 media were then prepared, and 30uL of these 10x solutions were directly pipetted into the wells containing the larvae, which have a total volume of ∼300uL, yielding 10uM Haloperidol, 1uM Pimozide, or 10uM Clozapine, in 0.1% DMSO vehicle. 30uL of 1% DMSO in E3 solution was pipetted into the vehicle control wells, yielding 0.1% DMSO vehicle control treated larvae. Larvae were treated with drug for between 30 and 90 minutes before the first dark flash was delivered. Vehicle control and drug treated larvae for each comparison were from the same clutches of larvae, and were assayed in different wells in the same behavioural plate.

### Data and software availability

C# and Matlab code for tracking and behavioural analyses, Arduino code for delivering stimuli, and laser cutting templates are available upon request.

## Acknowledgements

We thank the Granato, Engert and Schier lab members for helpful advice regarding the manuscript and work. This work was supported by the NIH grant RO1 MH092257 (M.G.), the NIH Brain Initiative grants U19NS104653, R24 NS086601 and R43OD024879, as well as a Simons Foundation grants SCGB# 542973 and 325207 (F.E).

